# The Shot CH1 domain recognises a distinct form of F-actin during *Drosophila* oocyte determination

**DOI:** 10.1101/2023.01.18.524359

**Authors:** D. Nashchekin, I. Squires, A. Prokop, D. St Johnston

**Affiliations:** The Gurdon Institute and the Department of Genetics, University of Cambridge; Tennis Court Road, Cambridge, United Kingdom; The University of Manchester, Manchester Academic Health Science Centre, Faculty of Biology, Medicine and Health, School of Biology, Manchester, United Kingdom

**Author notes:** Biosciences Institute, Newcastle University, Newcastle upon Tyne, United Kingdom (I. Squires).

## Abstract

As in mammals, only one cell in a *Drosophila* multicellular female germline cyst is specified as an oocyte. The symmetry-breaking cue for oocyte selection is provided by the fusome, a tubular structure connecting all cells in the cyst. The *Drosophila* spectraplakin Shot localises to the fusome and translates its asymmetry into a polarised microtubule network that is essential for oocyte specification, but how Shot recognises the fusome is unclear. Here we demonstrate that Shot’s actin-binding domain (ABD) is necessary and sufficient to localise Shot to the fusome and mediates Shot function in oocyte specification together with the microtubule-binding domains. The calponin homology domain 1 (CH1) of Shot’s ABD recognises fusomal F-actin and requires CH2 to distinguish it from other forms of F-actin in the cyst. By contrast, the ABDs of Utrophin, Fimbrin, Filamin, Lifeact and F-tractin do not recognise fusomal F-actin. We therefore propose that Shot propagates fusome asymmetry by recognising a specific conformational state of F-actin on the fusome.

## Introduction

Both male and female gametes differentiate inside cysts of interconnected germ cells. Whereas all male cells in the cyst become sperm, only one or few of the female germ cells are specified to become oocytes in most animals (Lu et al., 2017, Pepling and Spradling, 1998, Lei and Spradling, 2013). Since cells in the female germline cyst share cytoplasm through intercellular bridges, there must be specific mechanisms to select the future oocyte. In *Drosophila*, a polarised microtubule (MT) network extends throughout the cyst and directs the dynein-dependent transport of oocyte fate determinants into one cell (Theurkauf et al., 1993; Li et al., 1994). A similar mechanism could also be involved in oocyte specification in the mouse (Lei and Spradling, 2016; Niu and Spradling, 2022).

In *Drosophila*, cyst formation and oocyte specification occur in the germarium at the anterior of the fly ovary (Fig. 1A). Oocyte determination starts when a germline stem cell divides asymmetrically to produce a cyst progenitor, a cystoblast, that goes through 4 rounds of incomplete division to produce a cyst of 16 cells connected by intercellular bridges called ring canals (de Cuevas et al., 1997). The cystoblast contains a spherical structure inherited from the stem cell called the spectrosome, which contains endoplasmic reticulum, spectrins and actin-binding proteins (Lighthouse et al., 2008). At each subsequent division, new spectrosomal material forms in the ring canal connecting the two daughter cells and this fuses with the pre-existing spectrosome to form the fusome, which becomes a branched structure extending into all 16 cells of the cyst (Lin et al., 1994; De Cuevas and Spradling, 1998). Because one cell inherits the original spectrosome/fusome from the cystoblast, this cell contains more fusomal material than the others and this ultimately specifies it as the pro-oocyte (Lin and Spradling, 1995; De Cuevas and Spradling, 1998).

**Figure 1.**
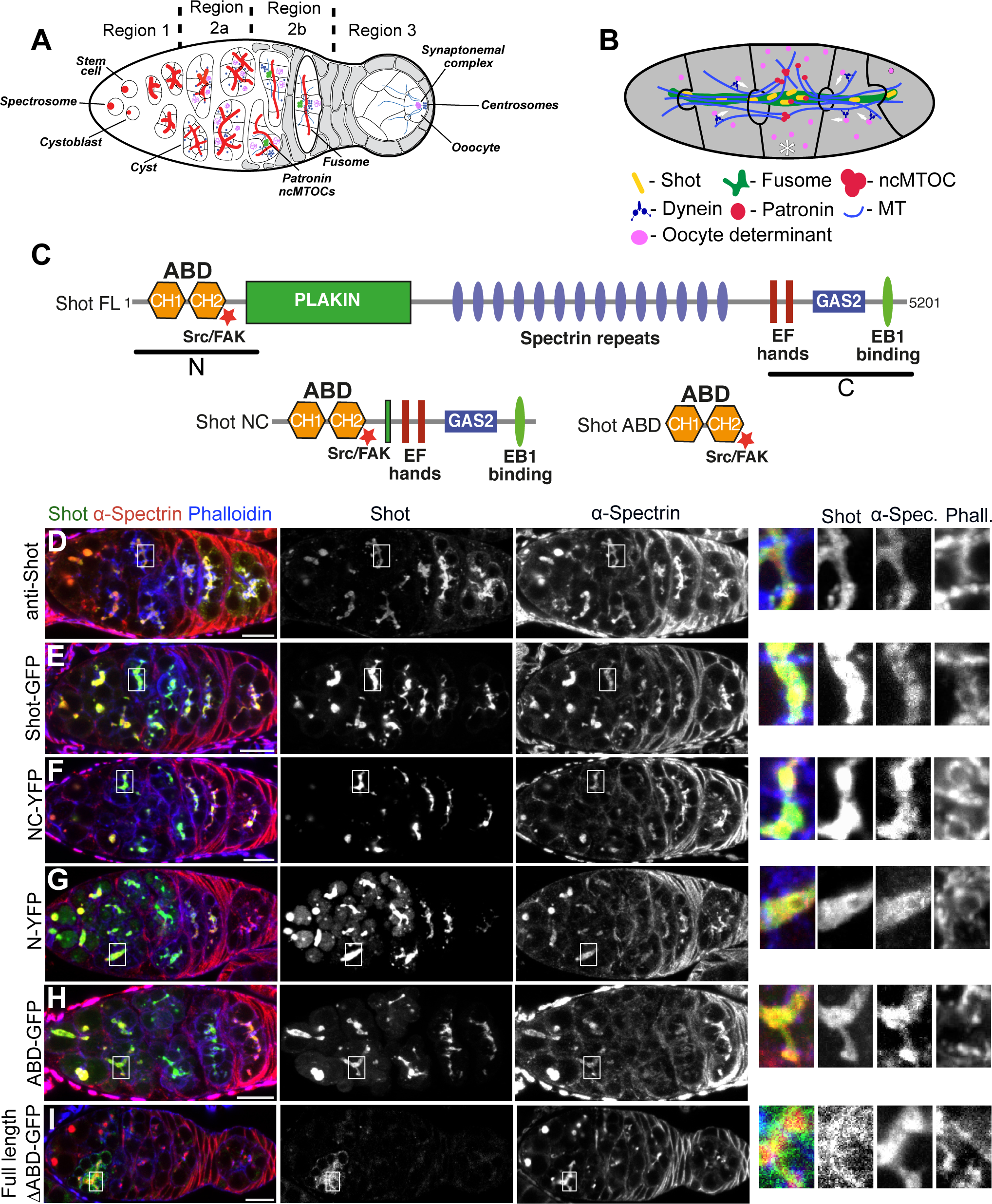
Shot is recruited to the fusome by its actin-binding domain. **(A)** Schematic diagram of the *Drosophila* germarium showing germline cyst formation and oocyte specification. **(B)** Diagram of a germline cyst showing how Shot and Patronin translate fusome asymmetry into the polarised MT network that directs dynein transport of oocyte fate determinates into the prospective oocyte (asterisk). Adapted from Nashchekin et al., 2021. **(C)** The domain structure of full-length Shot and Shot truncations. CH, calponin homology domain. ABD, actin-binding domain. **(D-I)** Endogenous Shot **(D)** and ectopically expressed full-length Shot-GFP **(E)**, Shot-NC-YFP **(F)**, Shot-N-YFP **(G)**, Shot ABD-GFP **(H)** localise to the fusome, whereas Shot^ΔABD^ **(I)** does not. An enlargement of the fusome is shown on the right. a-Spectrin marks fusome. Phalloidin marks ring canals and the cell cortex. Scale bars, 10µm.

The first step in the translation of fusome asymmetry into oocyte specification is the recruitment of the *Drosophila* spectraplakin, Shot (Roper and Brown, 2004; Nashchekin et al., 2021). Shot in turn recruits the MT minus end-binding protein Patronin (CAMSAP in mammals) to the fusome, where Patronin stabilises microtubule minus ends (Goodwin and Vale, 2010; Jiang et al., 2014; Nashchekin et al., 2021). The slight excess of Patronin in the future oocyte is then amplified by the dynein-dependent transport of Patronin and microtubule minus ends along the stabilised microtubules into this cell, leading to the formation of non-centrosomal microtubule-organising centres (ncMTOCs) in the future oocyte. Finally, these ncMTOCs nucleate a polarised MT network that directs the transport of oocyte determinants into this cell (Grieder et al., 2000; Bolívar et al., 2001; Nashchekin et al., 2021) (Fig. 1B).

The spectraplakin Shot belongs to a conserved family of actin-microtubule crosslinkers that includes human Dystonin and MACF1/ACF7, which play important roles in cytoskeletal organisation during neurogenesis and in epithelia (Lee and Kolodziej, 2002; Voelzmann et al., 2017; Dogterom and Koenderink, 2018). Spectraplakins are characterised by an N-terminal actin-binding domain (ABD), a central long rod domain consisting of plakin and spectrin repeats and a C-terminal MT-binding module (Fig. 1C). The ABD of spektraplakins consists of tandem calponin homology domains, CH1 and CH2 and the MT-binding module is composed of the MT lattice-binding GAS2 domain and an unstructured C-terminal domain containing two SxIP motifs that interact with the MT plus end-binding protein EB1 (Sun et al., 2001; Honnappa et al., 2009; Applewhite et al., 2010; Alves-Silva et al., 2012). Although Shot transmits fusome asymmetry to Patronin localisation and the formation of the polarised MT network that specifies the oocyte, how Shot recognises the fusome is not known. Previous work suggested that the Shot ABD is not involved (Roper and Brown, 2004). Here we show, however, that both the actin and MT-binding domains are required for oocyte specification, consistent with Shot’s role in organising the polarised microtubule network. The Shot ABD is necessary and sufficient for localisation to the fusome and recognises a form of F-actin that differs from other F-actin networks in the cyst.

## Results and Discussion

### The Shot actin-binding domain localises to the fusome

To determine which Shot domain(s) direct its localisation to the fusome, we expressed a mini-version of Shot lacking the central rod domain, Shot-NC (Figure 1C). It has been previously shown that the NC version of ACF7 (a mammalian Shot homologue) partially substitutes for ACF7 function in cells (Wu et al., 2008). Like endogenous Shot and the full-length Shot transgene (Fig. 1D and 1E, respectively), Shot-NC localised to the fusome, which is marked by a-spectrin (Fig. 1F and S1A). Thus, the fusome-binding activity of Shot resides in either its N- or C-terminal domains and the rod domain is dispensable for fusome localisation. Expression of the N- or C-terminal domains alone showed that Shot-N binds to the fusome in wild type and *shot* mutant cysts (Fig. 1G and Fig. 2C, respectively), whereas Shot-C forms cytoplasmic foci and accumulates in one cell of the cyst in wild-type (Fig. S1A and S1B) but not in *shot* mutant cysts (Fig. S1D), a pattern previously described for EB1 (Nashchekin et al., 2021). Live imaging of Shot-C-YFP in the germarium revealed that it forms EB1-like comets (Video1) suggesting that the C-terminal domain of Shot associates with MT plus ends, and that the MT lattice-binding GAS2 domain is in an inhibited conformation, as shown for the mammalian Shot homologue, Dystonin, in Cos-7 cells (Kapur et al., 2012). Nevertheless, we decided to test whether the GAS2 domain has the potential to bind the fusome by expressing a portion of the Shot C-terminal domain containing the EF hand and the GAS2 domain that has strong MT binding activity (Maybeck and Roper, 2009). EF-GAS2-GFP localised to the fusome in the presence of endogenous Shot (Fig. S1A and S1B) but failed to do so in *shot* mutant cysts (Fig. S1C and S1D). Fusome-associated MTs are largely lost in the absence of Shot, indicating that EF-GAS2-GFP localises to the fusome by binding to Shot-dependent MT (Roper and Brown, 2004). The GAS2 domain therefore cannot be responsible for the initial recruitment of Shot to the fusome.

**Figure 2.**
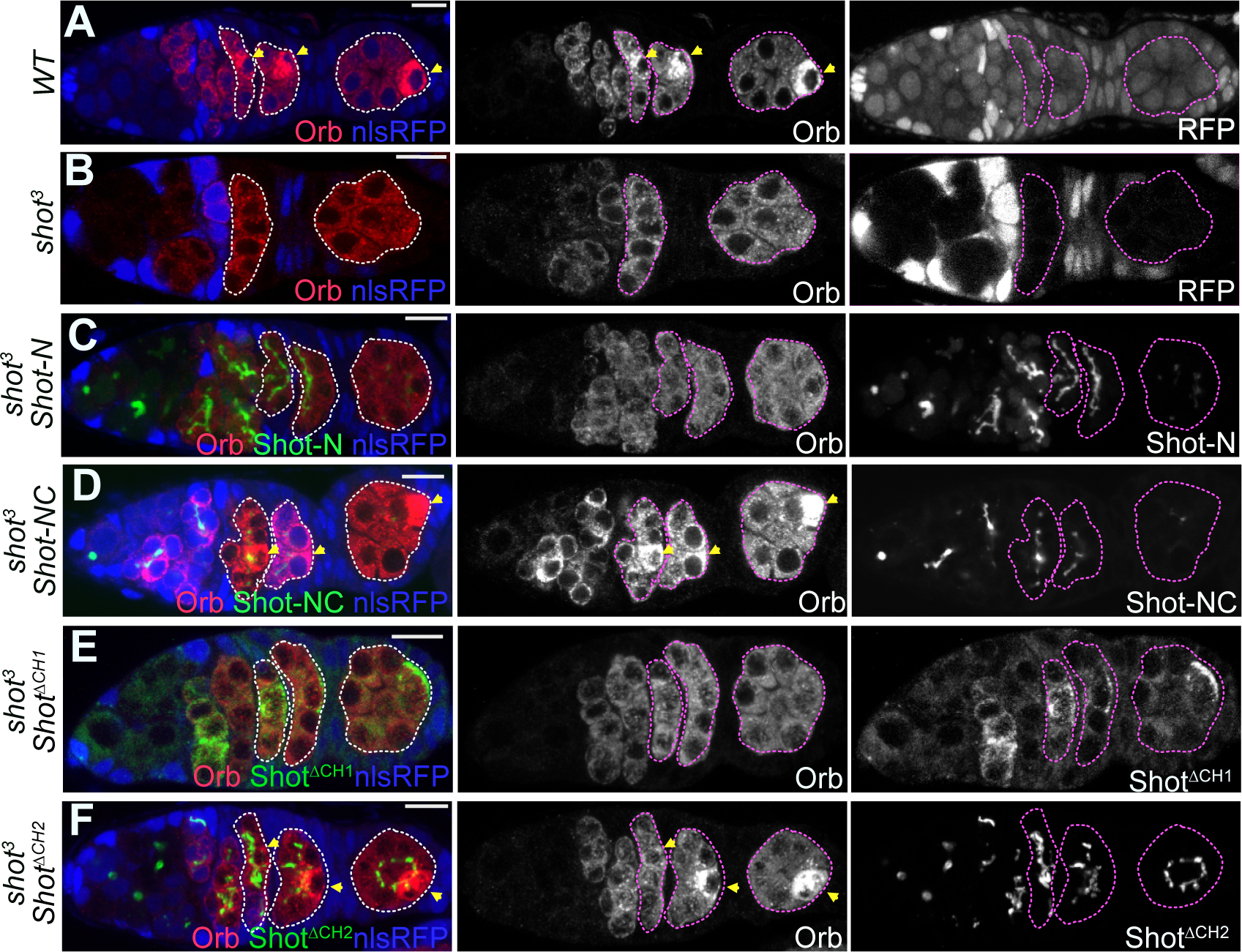
The Shot CH1 and MT-binding domains are required for the oocyte determination. **(A-B)** The distribution of Orb in wild type (WT) **(A)** and *shot^3^* **(B)** germline clone mutant cysts marked by the loss of nlsRFP (blue). **(C)** Shot-N-YFP does not rescue oocyte determination in *shot^3^* germline clones. **(D)** Expression of Shot NC-YFP in *shot^3^* mutant cysts restores oocyte specification. **(E-F)** Expression of Shot^ΔCH2^-GFP **(F)** but not Shot^ΔCH1^-GFP **(E)** rescues oocyte specification in *shot^3^* germline clones. Arrows point to the future oocyte; cysts are marked by dashed lines; mutant cysts are labelled by the absence of nuclear RFP (nlsRFP, blue). Scale bars, 10µm.

From our results so far, we speculated that Shot recruitment to the fusome requires the Shot N-terminal region containing the two Calponin homology domains that constitute the ABD. We therefore expressed a construct containing just the ABD of Shot fused to GFP and confirmed its localisation to the fusome (Fig. 1H and S1A). In contrast, full-length Shot lacking the ABD (Shot^ΔABD^) showed only a residual fusome localisation (Fig. 1I and S1A) that disappeared in the absence of endogenous Shot (Fig. S1D) like Shot C and Shot EF-GAS. Thus, the ABD of Shot is both necessary and sufficient for fusome localisation, presumably by interacting with fusome-associated F-actin.

### The actin- and MT-binding domains of Shot are required for the oocyte specification

The oocyte is not specified in the absence of Shot, leading to the formation of a 16 nurse cell follicle, but which domains of Shot are required for oocyte specification is not known (Roper and Brown, 2004). The cytoplasmic polyadenylation element binding factor oo18 RNA-binding protein (Orb) is an early marker for oocyte specification that becomes concentrated in the oocyte in regions 2b and 3 in wild-type germaria but it is uniformly distributed in *shot* null mutant cysts (Lantz et al., 1994; Roper and Brown, 2004) (Fig. 2A and 2B). To determine whether the presence of the Shot actin- or MT-binding domains are sufficient for oocyte specification, we expressed Shot-N, Shot-C, Shot-NC and Shot-EF-GAS2 in *shot* null mutant cysts (Fig. 2C-D and S1D). Although Shot-N bound to the fusome in the absence of endogenous Shot, it did not rescue Orb localisation and oocyte determination (Fig. 2C). Neither Shot-C or Shot-EF-GAS2 localised to the fusome or rescued oocyte specification in *shot* mutants (Fig. Fig. S1C-D). However, Shot-NC expression restored oocyte specification in the absence of full length Shot (Fig. 2D). Thus, both the ABD and MT-binding modules are necessary for Shot function during oocyte determination, suggesting that Shot acts as an actin-microtubule cross-linker in this context.

It has been previously reported that Shot’s ABD is not required for oocyte specification, since the oocyte is specified normally in cysts mutant for *shot^kakp1^,* a P-element insertion that is predicted to prevent the expression of CH1 domain-containing Shot isoforms (Roper and Brown 2004). As shown above, however, Shot’s ABD is essential for the Shot localisation to the fusome and oocyte determination. To resolve this contradiction, we tested the requirement for Shot’s CH1 and CH2 domains in oocyte specification by expressing Shot^ΔABD^, Shot^ΔABD^-LifeAct, Shot^ΔCH1^ and Shot^ΔCH2^ in *shot* mutant cysts (Fig. 2E-F and S1D). Shot truncations lacking the CH1 domain did not rescue oocyte specification, nor did substituting the Shot ABD with the actin-binding activity of LiveAct (Fig. 2E and S1D). Only *shot* mutant cysts expressing Shot^ΔCH2^ maintained Orb localisation and specified the oocyte (Fig. 2F). Thus, the CH1 domain of Shot’s ABD is essential for the oocyte specification. Since the *shot^kakp1^* mutant does not affect oocyte determination, we assume that this P-element insertion does not disrupt the expression of CH1-containing Shot isoforms in the germ line, although it does so in somatic tissues (Roper and Brown 2004).

Previously, it has been proposed that Shot binds the fusome with an unidentified domain and uses its GAS2 domain to bind and stabilise MT (Roper and Brown, 2004). Based on our results we propose an alternative model for Shot function in oocyte determination where it works as a classical actin-MT cytolinker by recognising fusomal F-actin with its ABD and using its C-terminal domain to attach MT to the fusome. Which part of the MT module is involved in this process is an open question. On the one hand, Shot-EF-GAS2 can recognise fusomal MT. On the other hand, expression of the whole Shot C-terminal domain showed that the GAS2 domain is not exposed and Shot C interacts only with MT plus ends through its EB1-binding motifs. Thus, Shot may guide the growth of MT plus ends along the fusome in a similar manner to that described for ACF7 in migrating cells (Kodama et al., 2003; Wu et al., 2008). The role of Shot in stabilising MT on the fusome could also be indirect, since we recently showed that Shot is required for the fusome localisation of the MT minus-end stabilising protein Patronin/CAMSAP (Nashchekin et al., 2021). Moreover, fusome-associated MT are unstable in *patronin* mutant cysts even though Shot is still present. How Shot’s C-terminus recruits microtubules to the fusome will therefore require further study.

### Shot recognises a distinctive form of F-actin on the fusome

Several actin-binding proteins have been identified as components of the fusome, including b-spectrin, Hts/Adducin, Tropomodulin and Shot (De Cuevas et al., 1996; Lin et al., 1994; Lighthouse et al., 2008; Roper and Brown, 2004). However, actin has never been detected in the fusome and Phalloidin does not label the fusome, staining only the ring canals and the cell cortex (Warn et al., 1985 and Fig. 1D-I). In contrast, Actin-GFP weakly localises to the fusome, raising the possibility that the absence of Phalloidin staining might be misleading (Fig. S2A). We therefore tested for the presence of endogenous actin in the fusome by performing antibody stainings for actin using various fixation methods. Only a combination of heat fixation followed by post-fixation with formaldehyde produced convincing fusomal actin staining (Fig. 3A). As a control we used *hts* mutant cysts (Fig. S2B), which lack the fusome (Lin et al., 1994), and confirmed that fusome-like actin staining was absent. We conclude that the fusome does contain F-actin, but not in a form that can be detected by Phalloidin. Since the Phalloidin-actin interaction is sensitive to the structure of F-actin filaments (McGough et al., 1997), fusomal F-actin may exist in a distinct conformation that can be bound by the Shot ABD but not by Phalloidin.

**Figure 3.**
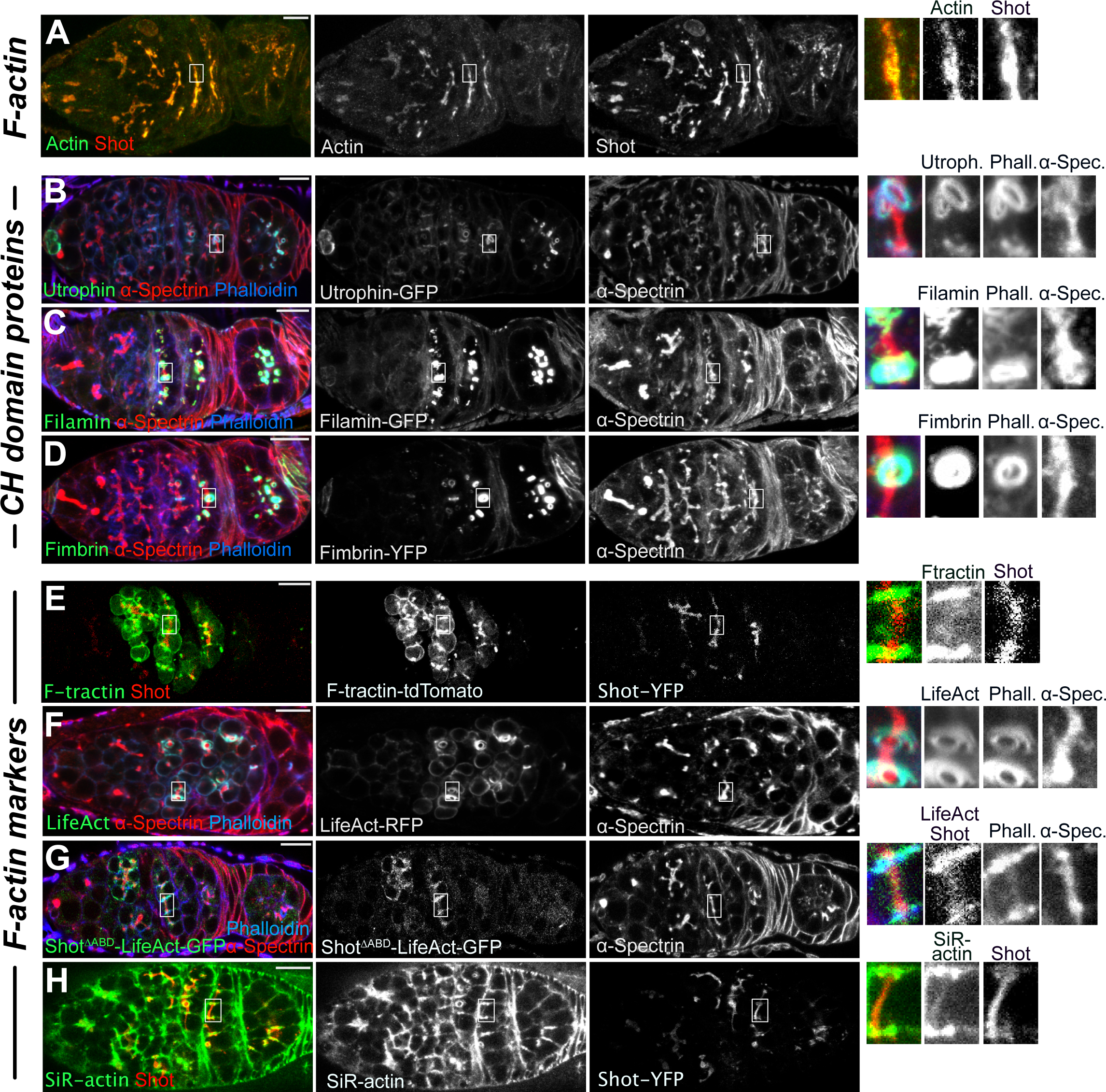
Standard F-actin labelling reagents and CH domain proteins fail to recognise fusomal F-actin. **(A)** Detection of F-actin on the fusome. A confocal image of a germarium stained with anti-actin after heat fixation and post-fixation with fomladehyde (green) and anti-Shot antibody (red). **(B-D)** Localisation of CH domain proteins (green) in cysts stained for a-spectrin (red) to label the fusome and cell cortices and Phalloidin (blue), which labels the ring canals. Utrophin ABD-GFP **(B)**, Filamin-GFP **(C)** and Fimbrin-YFP **(D)** localise to ring canals and the cell cortex but not the fusome. **(E)** A confocal image of a living germarium expressing F-tractin-tdTomato (green) and Shot-YFP (red). Shot localises to the fusome, whereas F-tractin labels the ring canals and cell cortex. **(F-G**) LifeAct does not recognise the fusome. Confocal images of germaria expressing LifeAct-RFP (green) **(F)** and Shot^ΔABD^-LifeAct-GFP stained for a-spectrin (red) and Phalloidin (blue) **(G). (H)** A confocal image of a living germarium expressing Shot-YFP (red) and incubated for 30 minutes with SiR-actin (green). A weak SiR-actin signal is present on the fusome. The righthand panels show enlargements of the fusome region. Scale bars, 10µm.

Both b-Spectrin and Shot are members of a large family of actin-binding proteins with ABDs formed by tandem CH domains (Korenbaum and Rivero, 2002; Yin et al., 2020) (Fig. S2C). To test whether other CH domain proteins recognise fusomal F-actin, we analysed the distribution of over-expressed Utrophin ABD and endogenously-tagged Filamin and Fimbrin. None of these CH domain proteins recognised fusomal F-actin and they mainly concentrated at the ring canals in fixed and live samples (Fig. 3B-D and S2D, respectively). F-tractin, LifeAct and SiR-actin are other commonly used F-actin markers (Melak et al., 2017), recognising a variety of F-actin structures (Riedl et al., 2008; Schell et al., 2001; Belin et al., 2014; Spracklen et al., 2014; D’Este et al., 2015). Neither F-tractin or LifeAct localised to the fusome, mainly localising to the cell cortex and ring canals (Fig. 3E-F and S2D). Moreover, substituting Shot’s ABD with the LifeAct sequence (Shot^ΔABD^-LifeAct) did not restore fusome recognition to full-length Shot (Fig. 3G and S1A). On the other hand, SiR-actin weakly localised to the fusome (Fig. 3H). Thus, fusomal F-actin has a distinctive conformation that is only recognised by a subset of actin-binding proteins/reagents. The Shot ABD must therefore have structural features that allow it to bind preferentially to fusomal F-actin and to distinguish it from other F-actin structures in the cyst. This does not preclude the Shot ABD binding to other forms of F-actin, since Shot re-localises to the cell cortex and ring canals in *hts* and **a*-spectrin* mutant cysts that lack the fusome (Lin and Spradling, 1995; De Cuevas et al., 1996) (Fig. S3). Shot also binds to actin filaments at the ring canals at the later stages of oogenesis when the fusome is disassembled (Lu et al., 2021).

The spatial arrangement of tandem CH1 and CH2 domains can regulate their actin binding activity, as the CH2 domain can sterically block some of the actin binding surfaces on the CH1 domain (Bañuelos et al., 1998; Galkin et al., 2010; Iwamoto et al., 2018). It has been proposed that phosphorylation of a conserved Tyr located at the end of the CH2 domain facilitates the formation of an “open” conformation of the ACF7 ABD and enhances F-actin binding (Yue et al., 2016, Fig. 1C). To test whether Shot binding to the fusome is regulated by Tyr phosphorylation, we expressed Shot ABDs containing phosphomimetic and non-phosphorylatable versions of Tyr364 (ABD Y364D and ABD Y364F, respectively). Whereas ABD Y364F only bound to fusomal F-actin (Fig. 4B), ABD Y364D bound to the fusome and ring canals in region 2a of the germarium and mostly relocalised to ring canals in region 2b (Fig. 4A). This suggests that shifting the equilibrium towards an open state of the CH domains alters the specificity of Shot’s ABD for different forms of F-actin, rather than working as a simple on/off switch.

**Figure 4.**
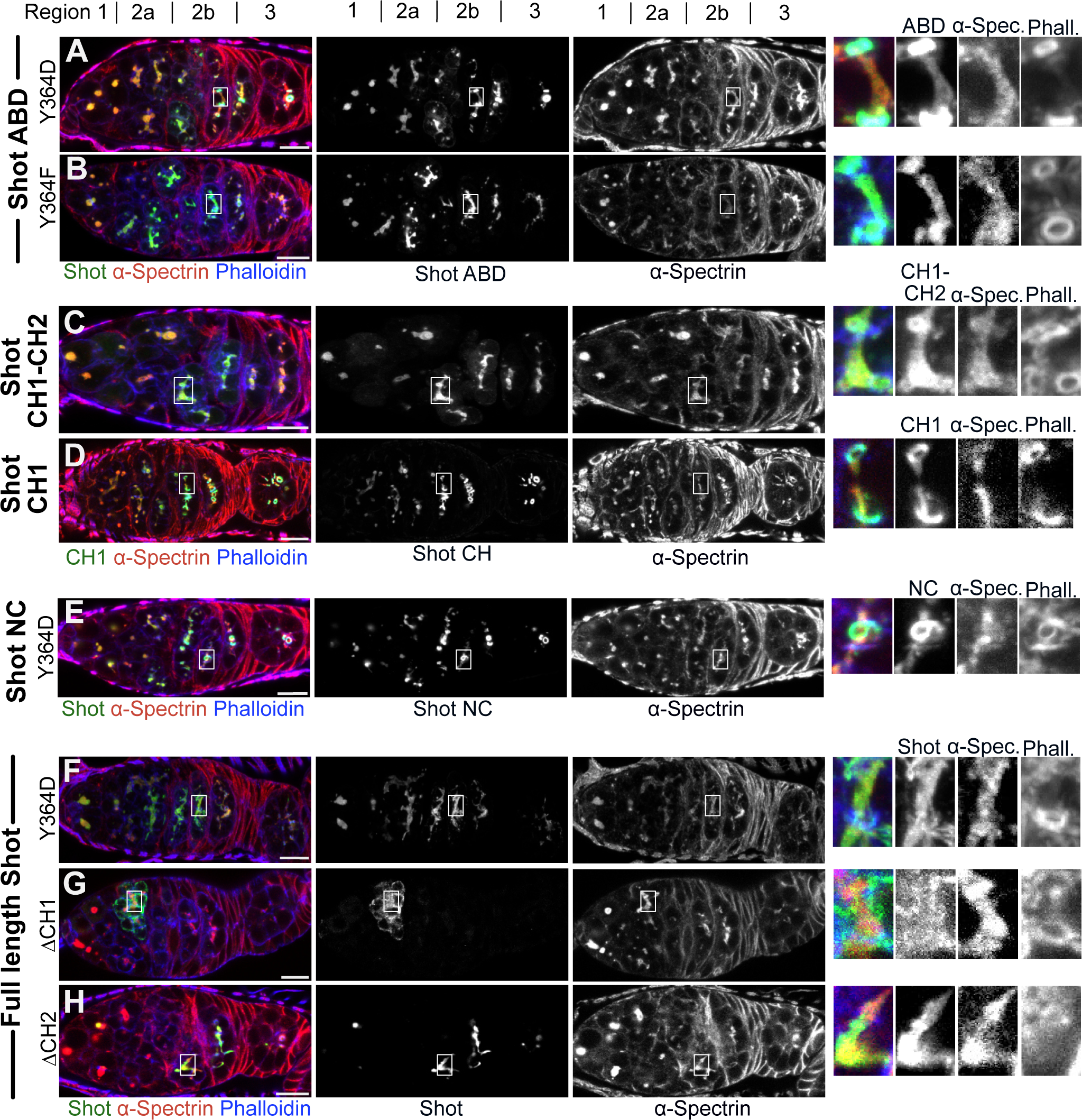
The Shot CH1 domain recognises the fusome. **(A-B)** The effects of phosphomimetic Y364D and non-phosphorylatable Y364F mutations. Y364D interferes with Shot ABD localisation to the fusome. Shot-GFP-ABDY364D **(A)** mostly localises to ring canals in region 2b of the germarium. Shot GFP-ABDY364F **(B)** localises like wild-type Shot. **(C-D)** Shot CH1-CH2 localises to the fusome whereas Shot CH1 localises to the fusome and ring canals. Germaria expressing Shot GFP-CH1-CH2 **(C)** and Shot GFP-CH1 **(D)**. **(E)** Shot-NCY364D-YFP mostly localises to ring canals in region 2b of the germarium. **(F)** A germarium expressing ShotY364D-GFP. Mutation of Y364 in full length Shot does not affect Shot localisation. **(G-H)** Deletion of the Shot CH1 domain abolishes binding to the fusome, but CH2 is not required. Germaria expressing Shot^ΔCH1^-GFP **(G)** and Shot^ΔCH2^-GFP **(H)**. The righthand panels show enlargements of the fusome region. a-Spectrin (red in the lefthand panels) marks the fusome. Phalloidin (blue) marks the ring canals and cell cortex. Regions of the germarium are indicated on the top. Scale bars, 10µm.

To examine the role of CH2 in regulating the actin-binding properties of CH1, we compared the localisation of CH1 alone with that of the full ABD containing CH1 and CH2. Whereas CH1-CH2 exclusively localised to the fusome (Fig. 1H and 4C), CH1 alone recognised F-actin on the fusome and ring canals (Fig. 4D). Thus, removing the CH2 domain has the same effect as the Y364D mutation in making CH1’s binding to F-actin more promiscuous. According to recent structural studies, CH1-CH2 binds to F-actin filaments in an open conformation, with CH1 sitting in the groove between subdomains 1 and 2 of an actin monomer and CH2 oriented away from the filament (Galkin et al., 2010; Iwamoto et al., 2018; Kumari et al., 2020). This implies that binding of CH1 to F-actin breaks multiple interactions between the CH1 and the CH2 (Galkin et al., 2010; Yue at al., 2016; Iwamoto et al., 2018; Harris et al., 2019; Kumari et al., 2020; Harris et al., 2020). We can envisage two models for how CH2 controls the specificity of CH1’s actin binding. CH2 in the closed conformation of the ABD may itself bind weakly to the specific form of actin on the fusome and thereby increase the avidity of the initial interaction of CH1-CH2 with fusomal actin. This increased avidity would then ensure that the Shot ABD binds preferentially to fusomal F-actin, which therefore outcompetes other F-actin in the cyst for Shot binding. Alternatively, CH2 may mask some actin binding surfaces of CH1 in the “closed” conformation of the ABD, allowing the latter to bind fusomal actin with higher affinity than other forms of actin. In another words, the presence of CH2 may increase the affinity of the ABD for fusomal actin or decrease its affinity to other forms of actin.

Since isolated ABDs could behave differently from the full-length protein, we introduced the Tyr364 mutations into Shot-NC and full-length Shot. Whereas the Shot-NC Tyr364 mutants behave like the corresponding ABD mutants, the localisation of the full-length Shot was not affected by the Y364D mutation (Fig. 4E, 4F and S2E). Thus, sequences in the rod domain somehow restore the specificity of the Y364D ABD for fusomal actin. We also tested the role each CH domain in the context of the full-length protein by analysing the localisation of full-length Shot lacking CH1 (Shot^ΔCH1^) or CH2 (Shot^ΔCH2^). Shot^DCH2^ localised to the fusome normally (Fig. 4H and S1A), whereas deletion of CH1 abolished fusome binding, leading to cytoplasmic localisation (Fig. 4G and S1A). The same pattern was observed in the absence of the endogenous Shot (Fig. 2F and 2E, respectively). These results indicate that the CH1 domain mediates Shot binding to the fusome and that it can recognise a specific F-actin conformation that is invisible to several other CH-domain proteins and actin-binding molecules. This result also implies that specific binding to fusomal F-actin by Shot, which depends on CH2 in the isolated ABD, does not require CH2 in the context of the full-length protein.

Structural studies on the interaction of CH domains with F-actin have revealed that CH1 interacts with two adjacent actin monomers and is sensitive to the torque/helicity of F-actin filaments (Hanein et al., 1998; Iwamoto et al., 2018; Kumari et al., 2020; Harris et al., 2020). Changes in the helical twist of F-actin filaments can be caused by mechanical tension, by interactions with actin-binding proteins (Harris et al., 2018; Jégou and Romet-Lemonne, 2020; Harris et al., 2020), or by the bending of F-actin filaments promoted by differences in the nucleotide states of actin monomers (Reynolds et al., 2022; Oosterheert et al., 2022). It is therefore possible that CH1 domains of different actin-binding proteins are predisposed to bind F-actin filaments with specific helical twists, which could explain their distinct, but partly overlapping localisation patterns (Washington and Knecht, 2008; Harris et al., 2020; Jégou and Romet-Lemonne, 2021).

Although the Y367D mutation and the removal of CH2 lead to promiscuous binding of the Shot ABD to both the fusome and other F-actin in the cyst, they have no effect on the specific localisation of full-length Shot to the fusome. This indicates that some region of the full-length protein can substitute for CH2 in the “closed” conformation of the ABD. This is most likely part of the rod domain since the Y367D mutation still causes promiscuous actin binding in the context of ShotNC (Fig. 4E). As proposed for CH2 in the context of the ABD alone, this rescue by the rod domain could be due to an interaction between the rod and CH1 that masks some actin binding surfaces to reduce its affinity for nonfusomal actin. Since the rod is very long, it is also possible that this effect is indirect and acts by allowing the intramolecular interaction between the C-terminal EF hands and CH1 to achieve the same effect (Applewhite et al., 2013). Alternatively, the rod domain could increase the avidity of Shot’s interaction with the fusome by binding to some other fusomal component. Indeed, Shot has been proposed to bind to a-Spectrin, which is enriched on the fusome (Khanal et al., 2016; Lin et al., 1994; De Cuevas and Spradling, 1998). Since full-length Shot^DABD^ does not localise to the fusome, the rod domain is unlikely function as a strong fusome-binding domain itself, but may contribute to avidity in a similar way to that proposed above for CH2.

Our evidence suggests that fusome asymmetry is propagated to MT organisation in the *Drosophila* female germline cyst by the formation of a distinct type of F-actin on the fusome, which then recruits the spectraplakin Shot. How fusomal F-actin is formed and the structural basis for its recognition by actin-binding proteins remain to be determined. Considering that Shot is a conserved actin binding protein, it is possible that a similar mechanism is used in other contexts where F-actin filaments in a specific mechanical state are recognised by only a subset of actin-binding proteins.

## Material and methods

### Drosophila stocks

The following previously described *Drosophila melanogaster* mutant alleles and transgenic lines were used: FRTG13 *shot^3^* (Roper and Brown, 2004), *hts^1^* and *hts^01103^* (Yue and Spradling, 1992), **a*-spectrin^e2-26^* (Hülsmeier et al., 2007), UAS-GFP-Actin42A (Roper et al., 2005), UAS-GFP-Shot EF-GAS2 (Maybeck and Roper, 2009); UAS-Shot^ΔABD^-GFP, UAS-Shot^ΔCH1^-GFP, UAS-Shot^ΔCH2^-GFP and UAS-Shot^ΔABD^-LifeAct-GFP (Qu et al., 2022), Shot-YFP (Nashchekin et al., 2016), UAS-F-tractin-tdTomato (Spracklen et al., 2014),Filamin-GFP trap line (gift from K. Röper, MRC-LMB, UK), Fimbrin-YFP (Cambridge Protein Trap Insertion line 100066, (Lowe et al., 2014), sqh>Utrophin ABD-GFP (Rauzi et al., 2010). UAS-Shot-GFP, UAS-ShotY364F-GFP, UAS-ShotY364D-GFP, UAS-Shot-NC-YFP, UAS-Shot-NCY364D-YFP, UAS-Shot-NCY364F-YFP, UAS-Shot-N-YFP, UAS-Shot-C-YFP, UAS-GFP-Shot ABD, UAS-GFP-Shot ABDY364F, UAS-GFP-Shot ABDY364D, UAS-LifeAct-RFP were generated for this study.

### Drosophila genetics

Germline clones of *shot^3^ and *a*-spectrin^e2-26^*were induced by incubating larvae at 37^◦^ for two hours per day over a period of three days. Clones were generated with FRT G13 nlsRFP and FRT 2A nlsRFP, (Bloomington Stock Center) using the heat shock Flp/FRT system (Chou and Perrimon, 1992). Germline expression of UAS transgenes was induced by nanos>Gal4. All transgenes were expressed in a wild-type background unless otherwise specified.

### Molecular Biology

To generate pUASP Shot-GFP, three fragments of the *shot* RE cDNA were amplified from pUAST Shot-GFP (Lee and Kolodziej, 2002) and cloned together with EGFP into the pUASPattb vector. pUASPattb-Shot-NC-YFP was generated by PCR amplifying fragments from pUAST Shot-GFP corresponding to the first 520 aa (Shot-N) and last 462 aa (Shot-C) of Shot PE, cloning them together into pUASP-YFP-Cterm and then re-cloning into pUASPattb. pUASPattb-Shot-N-YFP and pUASPattb-Shot-C-YFP were generated by amplifying Shot-N or Shot-C fragments from pUASPattb-Shot-NC-YFP and cloning them together with YFP into pUASPattb. Shot ABD cDNA (corresponding to 146-368 aa of Shot PE) was amplified from pUASPattb-Shot-NC-YFP and cloned together with EGFP into pUASP-attb to generate pUASP-attb GFP-Shot ABD. The Q5 Site-Directed Mutagenesis Kit (New England BioLabs) was used to generate pUASPattb-GFP-Shot ABDY364F, pUASPattb-GFP-Shot ABDY364D, pUASPattb-Shot-NCY364D-YFP and pUASPattb-Shot-NCY364F-YFP. Shot-N with the Y364F or Y364D mutations was amplified from pUASPattb-Shot-NCY364F or pUASPattb-Shot-NCY364D, respectively, and cloned together with two fragments covering the rest of Shot RE cDNA and EGFP into pUASP-attb to generate pUASPattb-ShotY364F-GFP and pUASPattb-ShotY364D-GFP. pUASP-LifeAct-tagRFP was generated according to Riedl et al (2008). NEBuilder HiFi DNA Assembly (New England BioLabs) was used for most of the cloning. Primer sequences are available on request.

### Immunohistochemistry

Ovaries were fixed for 20 min at room temperature in 4% paraformaldehyde and 0.2% Tween in PBS. Ovaries were then blocked with 1% BSA in PBS with 0.2% Tween for 1 hr at room temperature. Ovaries were incubated with the primary antibody for 16 hr with 0.1% BSA in PBS with 0.2% Tween at 4C° and for 4 hr with the secondary antibody at room temperature. For detection of fusomal F-actin, we used the heat fixation protocol described in Chen et al (2018) with some modifications. Ovaries were dissected in PBS, fixed for 10 sec in hot (95C°) 1xTSS (0.03% Triton X-100, 4 g/L NaCl), post-fixed in 8% paraformaldehyde and 0.1% Triton X-100 in PBS for 10 min at room temperature and then treated as above. We used the following primary antibodies: guinea pig anti-Shot at 1:1000 (Nashchekin et al., 2016), mouse anti-Orb at 1:10 (DSHB Hybridoma Products 4H8 and 6H4. Deposited to the DSHB by Schedl, P), mouse anti-a-Spectrin at 1:200 (DSHB Hybridoma Product 3A9. Deposited to the DSHB by Branton, D. / Dubreuil, R.), mouse anti-Actin at 1:200 (clone AC-40, Merck), rabbit anti-b-spectrin at 1:200 (Byers et al., 1989), SiR-actin (1:100, Spirochrome). Secondary antibodies conjugated with Alexa fluor dyes (Thermo Fisher Scientific) were used at 1:1000.

### Imaging

Fixed preparations were imaged using a Leica SP8 (63x/1.4 HC PL Apo CS Oil) confocal microscope. Germaria were imaged by collecting 10–15 z sections spaced 0.5 µm apart. For live imaging, ovaries were dissected and imaged in Voltalef oil 10S (VWR International) on a Leica SP5 confocal microscope (63x/1.4 HCX PL Apo CS Oil) or on an Olympus IX81 inverted microscope with a Yokogawa CSU22 spinning disk confocal imaging system (100x/ 1.3 NA Oil UPlanSApo). Images were collected with Leica LAS AF software or MetaMorph and processed using ImageJ. The images are projections of several z sections.

### Reproducibility of experiments and statistical analyses

Images are representative examples from at least three independent repeats for each experiment. The number of *shot^3^*mutant cysts (region 2b to 3) with restored Orb localisation after the expression of a rescue transgene were as follows: Figure 2A wild type cysts (50/50), Figure 2B *shot^3^*mutant cysts without a transgene (0/30), Figure 2C Shot N (0/21), Figure S1D Shot-C (0/28), Figure S1D Shot EFGAS (0/18), Figure 2D Shot NC (25/26), Figure S1D Shot^ΔABD^ (0/14), Figure S1D Shot-LifeAct (0/19), Figure 2E Shot^ΔCH1^ (0/16), Figure 2F Shot^ΔCH2^ (21/21). The number of cysts analysed to determine the localisation of various Shot transgenes and actin-binding reagents is summarised in Table S1. The chi-square test was used to test whether values were significantly different between wild type and *shot^3^* mutant cysts. The Mann-Whitney t-test was used to determine significance when measuring the co-localisation of Shot transgenes and fusome. A level of p< 0.01 was considered to be statistically significant. No statistical methods were used to predetermine sample size, the experiments were not randomized, and the investigators were not blinded to allocation during experiments and outcome assessment.

## Supplemental material

Figures S1-S3, Video 1 and Table S1.

## Acknowledgements

We are grateful to Jean-René Huynh, Christian Klämbt, Thomas Lecuit, Katja Röper and the Bloomington Drosophila Stock Center (NIH P40OD018537) for fly stocks, the Gurdon Institute Imaging Facility for assistance with microscopy, John Overton for technical assistance in making transgenic flies. We thank Jia Chen for help with heat fixation.

## Figure legends

**Figure S1.**
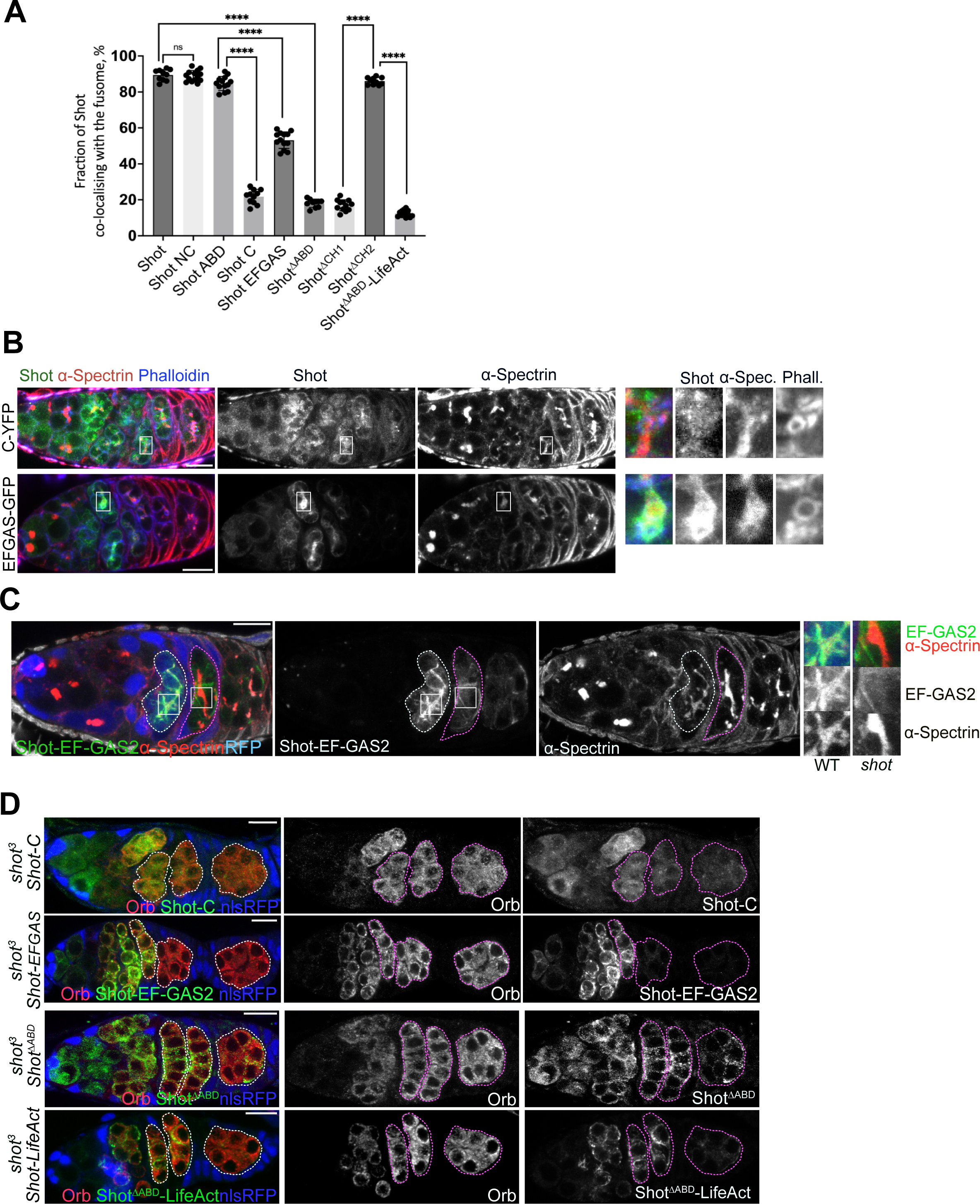
**(A)** Quantification of fusome and Shot co-localisation in germaria expressing various Shot transgenes and stained with anti-a-Spectrin to label the fusome. Manders’ co-localisation coefficient is measured using the JACoP plug-in for Fĳi (Bolte and Cordelieres, 2006). Data are means with SD. ****, P < 0.0001. **(B)** Ectopically expressed Shot-EF-GAS2-GFP localises to the fusome, whereas Shot C-YFP does not. An enlargement of the fusome is shown on the right. a-Spectrin marks the fusome. Phalloidin marks ring canals and the cell cortex**. (C)** Shot-EF-GAS2-GFP does not localise to the fusome in the absence of endogenous Shot. A germarium expressing Shot-EF-GAS2-GFP (green) in wild type and *shot*^3^ mutant cysts. An enlarged view of the fusome is shown on the right. a-Spectrin (red in left panel) marks the fusome. **(D)** Shot-C-YFP, Shot-EF-GAS2-GFP, Shot^ΔABD^-GFP and Shot^ΔABD^-LifeAct-GFP do not rescue oocyte determination in *shot^3^* germline clones. Cysts are marked by dashed lines; mutant cysts are labelled by the absence of nuclear RFP (nlsRFP; blue). Scale bars, 10µm.

**Figure S2.**
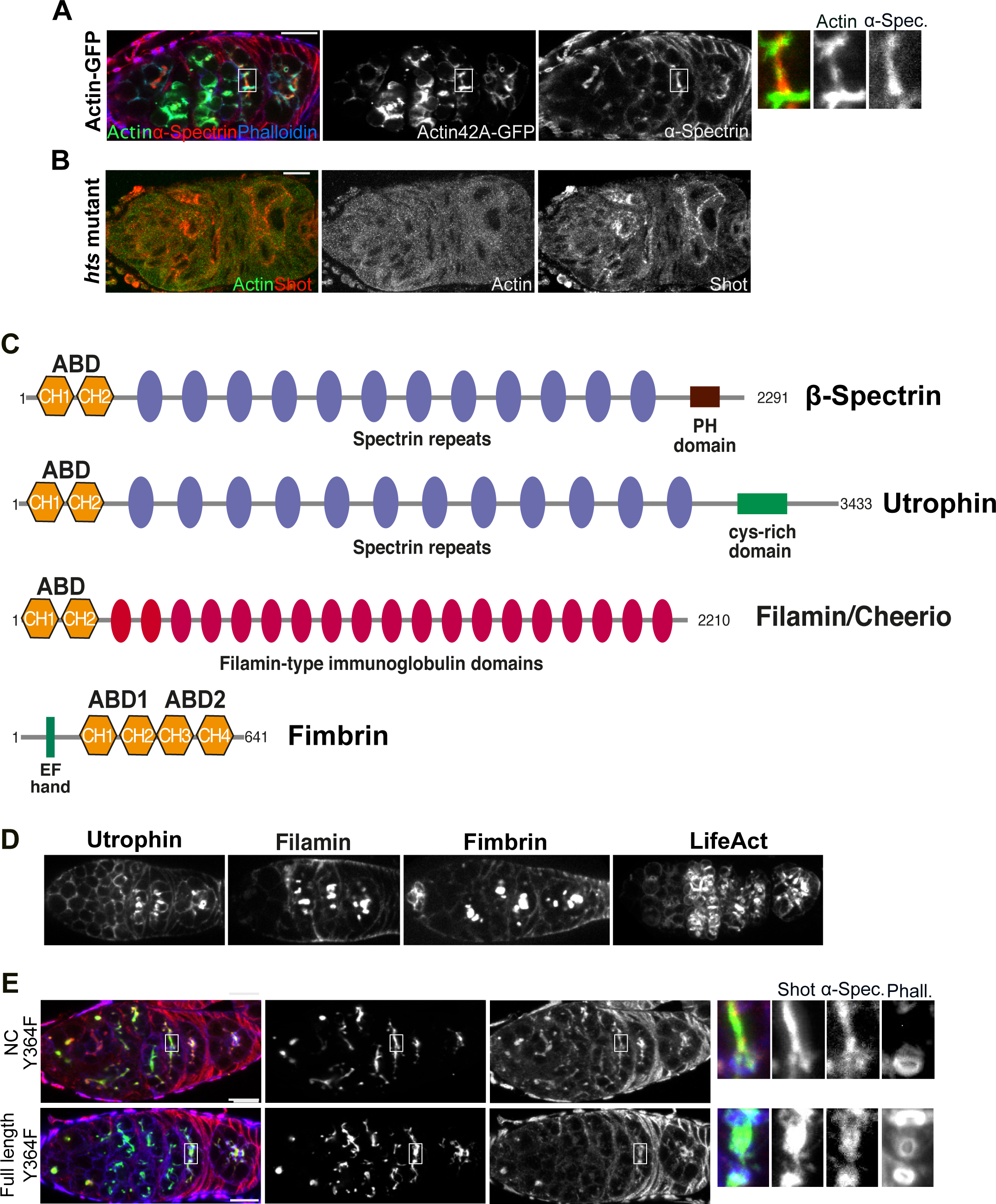
**(A)** Actin-42A-GFP localises to the fusome. A germarium expressing Actin-42A-GFP (green) stained with anti-a-Spectrin antibody (red) to mark the fusome. The righthand panel shows an enlarged view of the fusome. **(B)** Fusomal actin is not detected in *hts* mutant cysts. A germarium from a *hts* mutant female stained with anti-Shot (red) and anti-Actin (green) antibodies after heat fixation followed by post-fixation in 8% formaldehyde. **(C)** Diagrams showing the domain structure of actin-binding proteins with calponin homology (CH) domains. ABD, actin-binding domain. **(D)** Live germaria expressing Untrophin ABD-GFP, Filamin-GFP, Fimbrin-YFP and LifeAct-RFP. **(E)** The non-phosphorylatable mutation in Shot Tyr364 does not affect Shot NC and full length Shot localisation. Germaria expressing Shot-NC Y364F-YFP (top) and Shot Y364F-GFP (bottom). The righthand panels show enlargements of the fusome region. a-Spectrin (red in the lefthand panels) marks the fusome. Phalloidin (blue) marks the ring canals and cell cortex. Scale bars, 10µm.

**Figure S3.**
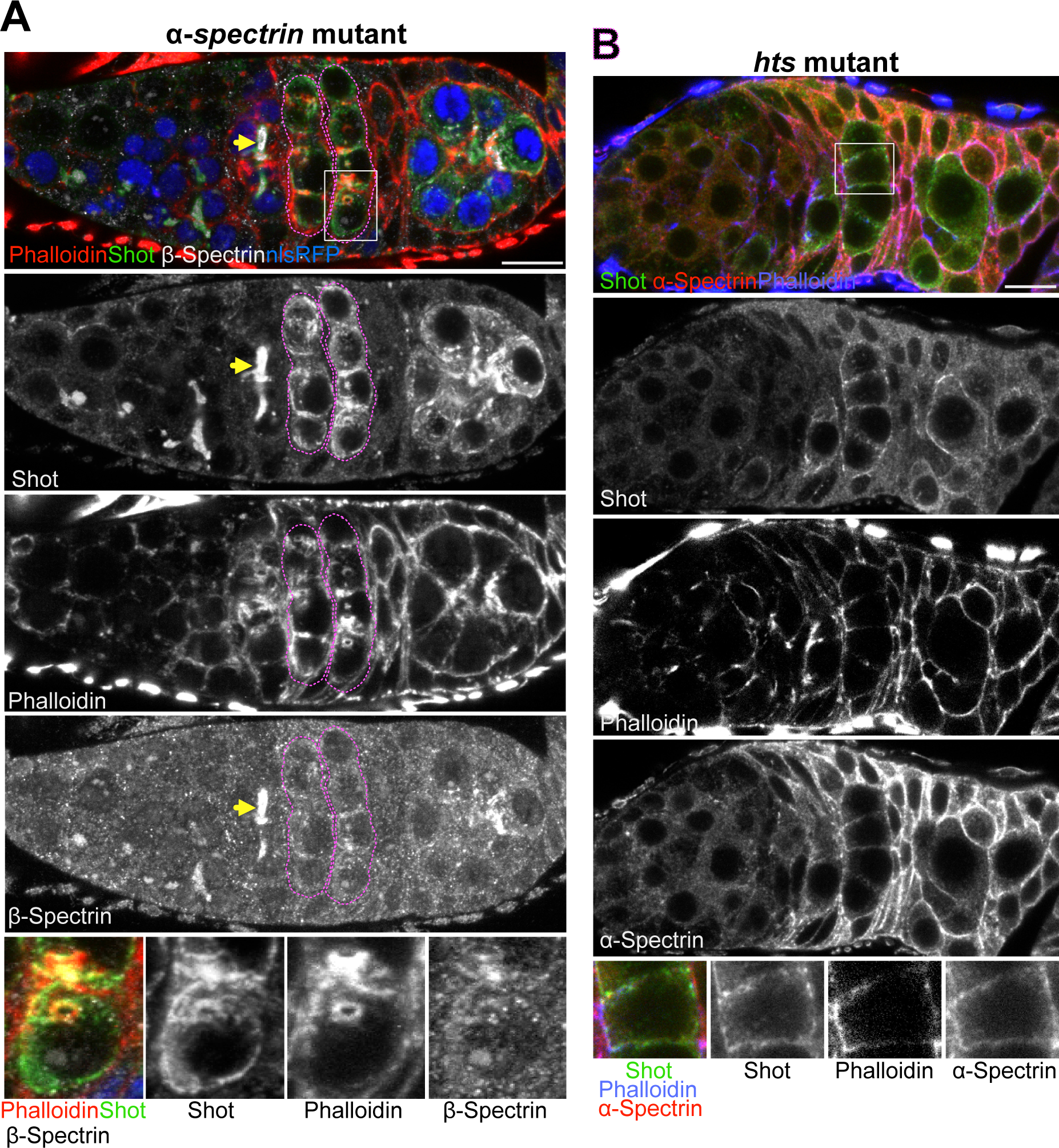
Shot localisation in *a-spectrin* and *hts* mutant cysts. Shot localises to the cell cortex and ring canals in *a-spectrin* **(A)** and *hts* **(B)** mutant cysts, which lack the fusome. **(A)** A germarium with *a-spectrin* germline clones stained with anti-Shot (red), Phalloidin (blue) and anti-b-Spectrin (white) to label the fusome. Mutant cysts are labelled by the absence of nuclear RFP (nlsRFP; blue). *a-spectrin* mutant cysts are marked by dashed lines. Arrows point to the fusome in a wild type cyst. **(B)** A germarium from a *hts* mutant female stained with anti-Shot (red), Phalloidin (blue) and anti-a-Spectrin (red) to label the fusome. Scale bars, 10µm.

## Video legends

**Video 1.** A time-lapse video of Shot-C-YFP in a wild-type germarium. Related to Figure S1B. Images were collected every 1 second on a spinning disc confocal microscope. The video is shown at 15 frames/sec.

**Table S1.** Localisation to the fusome and ring canals.

| Name of the protein | Localisation to the fusome / number of cysts analysed | Localisation to ring canals / number of cysts analysed | Number of germaria analysed |
| --- | --- | --- | --- |
| Shot FL | 63/63 | 0/63 | 11 |
| Shot NC | 71/71 | 0/71 | 10 |
| Shot N | 66/66 | 0/66 | 10 |
| Shot C | 0/42 | 0/42 | 10 |
| Shot EF-GAS2 | 32/32 | 0/32 | 8 |
| Shot ABD | 58/58 | 0/58 | 10 |
| Shot<sup>ΔABD</sup> | 0/34 | 0/34 | 16 |
| Shot<sup>ΔABD</sup>-LifeAct* | 0/41 | 5 /41 | 12 |
| Shot<sup>ΔCH1</sup> | 0/31 | 0/31 | 10 |
| Shot<sup>ΔCH2</sup> | 47/47 | 0/47 | 10 |
| Shot Y364D | 74/74 | 0/40 (region 2a)<br>0/34 (region 2b) | 10 |
| Shot Y364F | 45/45 | 0/25 (region 2a)<br>0/20 (region 2b) | 10 |
| Shot NC Y364D | 49/49 | 10/27(region 2a)<br>22/22(region 2b) | 10 |
| Shot NC Y364F | 45/45 | 0/24 (region 2a)<br>0/21(region2b) | 10 |
| Shot ABD Y364D | 55/55 | 14/31(region 2a)<br>24/24(region2b) | 10 |
| Shot ABD Y364F | 53/53 | 0/28 (region 2a)<br>0/25 (region 2b) | 10 |
| Shot CH1 | 53/53 | 13/29 (region 2a)<br>24/24 (region 2b) | 10 |
| Utrophin ABD | 0/50 | 50/50 | 10 |
| Filamin | 0/51 | 51/51 | 10 |
| Fimbrin | 0/42 | 42/42 | 10 |
| F-tractin | 0/49 | 49/49 | 12 |
| LifeAct | 0/42 | 42/42 | 8 |
| SiR-actin | 41/41 | 41/41 | 8 |

